# A Pan-Beta-Coronavirus Vaccine Bearing Conserved and Asymptomatic B- and T-Cell Epitopes Protect Against Highly Pathogenic Delta and Highly Transmissible Omicron SARS-CoV-2 Variants of Concern

**DOI:** 10.1101/2025.04.13.648648

**Authors:** Hawa Vahed, Swayam Prakash, Afshana Quadiri, Izabela Coimbra Ibraim, Etinosa Omorogieva, Swena Patel, Jimmy Tadros, Emma Jane Liao, Lauren Lau, Aziz A Chentoufi, Anthony B Nesburn, Baruch D Kuppermann, Jeffrey B. Ulmer, Daniel Gil, Lbachir BenMohamed

## Abstract

SARS CoV-2 continues to evolve into new viral variants due to mutation, primarily in the Spike protein. Existing Spike-based vaccines are less effective because these variants can be more transmissible and evade vaccine-induced immunity. By targeting more conserved, Spike, and non-Spike, viral antigens using both arms of the adaptive immune system, i.e. B and T cells, we aim to reduce the reliance on neutralizing antibodies and avoid potential mismatches between the COVID-19 vaccines and circulating virus strains. In this way, enhanced immune memory function and broad-spectrum protection against existing and evolving virus variants can be attained. We have developed a mRNA-LNP-based multi-epitope vaccine incorporating conserved CD8^+^ T-cell, CD4^+^ T-cell, and B-cell epitopes. These conserved epitopes were selected as being highly recognized by B- and T-cells from unvaccinated asymptomatic COVID-19 patients. To evaluate the effectiveness of this multi-epitope “asymptomatic” vaccine, we utilized a novel triple transgenic h-ACE-2-HLA-A2/DR mouse model to enable the assessment of human T cell epitopes. Key observations include induction of: (*i*) robust protection against infection and disease caused by SARS-CoV-2 Delta (B.1.617.2) and Omicron (XBB.1.5) variants, as measured by reduced weight loss, virus replication, and lung pathology; (*ii*) strong antibody responses,; and (*iii*) potent SARS-CoV-2 epitope-specific IFN-γ-producing CD4^+^/CD8^+^ T cells and Follicular helper CD4^+^ T (T_FH_) cells. These data support the strategy of targeting B-cells and T-cells directed toward highly conserved and “asymptomatic” epitopes, from both structural and non-structural viral protein antigens, to generate a broad-spectrum protective immunity to minimize disease impact across multiple SARS-CoV-2 variants.

## INTRODUCTION

The coronavirus disease 2019 (COVID-19) pandemic caused by SARS-CoV-2 has had a major impact on public health and economies globally. ^1^ SARS CoV-2 continues to circulate widely and mutate to generate viral variants. As of late January 2025, the SARS-CoV-2 Omicron variants belonging to lineages JN.1 (BA.2.86), LP.8.1 (10%), KP.3 (JN.1.11.1), KP.3.1.1 (JN.1.11.1), MC.1 (KP.3.1.1), and XEC (JN.1.13) have predominated in the United States. ^2^ These new variants often exhibit increased transmissibility and immune evasion, thereby resulting in reduced effectiveness of existing vaccines. Hence, although COVID-19 vaccines have played a pivotal role in reducing overall morbidity and mortality, significant challenges persist. ^3^ This is particularly true in the elderly and individuals with weakened immune systems or underlying comorbidities, who remain at high risk for serious disease outcomes. Initial Phase 3 clinical data in late 2020, showed remarkable levels of vaccine efficacy >90% for prevention of COVID-19 ^4^ ^5^. However, recent studies have shown that the 2023-2024 vaccine formulations provided modest protection against the SARS-CoV-2 JN.1 lineage ^6^ and little to none against the FLiRT (KP.2 & JN.1) & FLuQE-like circulating emerging SARS-CoV-2 variants, ^7^ likely as a result of a mismatch between the vaccine and circulating virus strains leading to reduced levels of neutralizing immunity against emerging Omicron subvariants. ^8^ These observations highlight the need for continuous monitoring and rapid adaptation of the vaccine to match circulating viral strains. ^9^ Or preferably to discover, develop, and implement a superior vaccine able to provide cross-strain protection.

Our strategy aims to minimize the reliance on neutralizing antibodies and the potential for a mismatch by targeting both arms of the adaptive immune system, including T and B cell responses to enhance immune memory formation and broader spectrum protection against existing and evolving virus variants. We have designed a multi-epitope vaccine comprising CD4^+^, CD8^+,^ and B cell epitopes conserved across diverse Coronavirus strains. Specifically, we used a two-pronged approach to identify and select our vaccine candidate. First, we employed an immuno-informatics approach to identify a total of 16 common CD8^+^ T cell epitopes, 6 common CD4^+^ T cell epitopes, and 8 potential B cell (antibody) epitopes that are highly conserved between 3.6 million SARS-CoV-2 strains that presently circulate in the United States and 200 other countries; and 20 variants including each of the SARS-CoV-2 alpha (α), beta (β), gamma (γ), delta (δ), and omicron (o) variants. Second, we selected human T cell epitopes that correlated with protection in unvaccinated, asymptomatic COVID-19 patients, as we have previously reported. ^10^ In this study, we utilized the established mRNA-LNP technology to deliver the antigenic epitopes and demonstrate safety, immunogenicity, and protective efficacy against the highly pathogenic Delta (B.1.617.2), and highly transmissible Omicron (XBB.1.5) SARS-CoV-2 variants in the novel triple transgenic HLA-A*02:01/HLA-DRB1*01:01-hACE-2 mice. ^10^

Taken together, these data provide preclinical proof of principle for a broad-spectrum vaccine bearing conserved and “asymptomatic” B, CD4^+^, and CD8^+^ T cell epitopes capable of providing cross-strain protection with the potential to reduce the need for frequent vaccine updating.

## MATERIALS & METHODS

### Viruses

We have used the highly pathogenic SARS-CoV-2 Delta (B.1.617.2) variant and the highly transmissible SARS-CoV-2 Omicron (XBB.1.5) variant in this experiment. The two SARS-CoV-2 variants were procured from Microbiologics (St. Cloud, MN). Upon receipt, in the BSL3 facility, we have propagated the SARS-CoV-2 variants Delta (isolate h-CoV-19/USA/MA29189; Batch number: G87167) and Omicron (isolate h-CoV-19/USA/FL17829; Batch number: G76172), to generate high-titer virus stocks in Vero E6 (ATCC-CRL1586). ^11^

### Sequence homology analysis among SARS-CoV-2 and animal CoV strains

We have used ∼8.5 million animal and human SARS-CoV-2 genome sequences for sequence homology analysis. These sequences were obtained from GISAID (www.gisaid.org) and NCBI databases. As described earlier ^10, 12^ the sequence homology analysis was performed to identify the highly conserved protein sequence regions across (*i*) all SARS-CoV-2 variants, (*ii*) common-cold SARS-CoV, (*iii*) MERS and SARS-CoV-1 strains, and (*iv*) animal CoV strains. For this sequence alignment purpose, we have used among others the DIAMOND ^13^ tool.

### Immunodominant CD8^+^ and CD4^+^ T Cell Epitopes prediction

For CD8^+^ and CD4^+^ T cell-based epitope prediction, we have used prediction tools like SYFPEITHI, MHC-I binding predictions, MHC-II Binding Predictions, Tepitool, and TEPITOPEpan were used. Sixteen 9-mer CD8^+^ T cell immunodominant epitopes and six 15-mer CD4^+^ T cell epitopes immunodominant epitopes were identified as we have reported earlier ^10, 12, 14^. Subsequently, peptides of >95% purity were synthesized for these immunodominant epitopes (21^st^ Century Biochemicals, Inc, Marlborough, MA).

### SARS-CoV-2 B Cell Epitope Prediction

Linear B cell epitope prediction was carried out on the spike glycoprotein (S) using the BepiPred 2.0 algorithm in the IEDB platform. A total of eight B cell peptides were obtained representing all the major non-synonymous mutations reported among the SARS-CoV-2 variants. Structure-based antibody prediction was performed using Discotope 2.0 as described before. ^12^

### mRNA synthesis and LNP formulation

mRNA encoding the multi-epitope vaccine was synthesized by *in vitro* transcription using T7 RNA polymerase (MegaScript, Thermo Fisher Scientific, Waltham, MA) on linearized plasmid templates, as previously reported. ^15^ Different linkers namely AAY (for CD8^+^ T cell epitopes), GPGPG (for CD4^+^ T cell epitopes), and KK (for B cell epitopes) were used in this particular vaccine construct (**Fig. 1B**). Modified mRNA transcript with full substitution of Pseudo-U was synthesized by TriLink Biotechnologies using proprietary CleanCap® technology. Purified mRNAs were analyzed by agarose gel electrophoresis and stored frozen at −20°C. The mRNAs were formulated into GenVoy-ILM-based LNPs using NanoAssemblr Ignite by Cytiva. mRNA LNPs were filtered and then analyzed for polydispersity and dynamic light scattering. RNA encapsulation efficiency and concentrations were determined using a RiboGreen plate-based assay. Formulated mRNA-LNPs were stored at −80°C.

**Figure 1.**
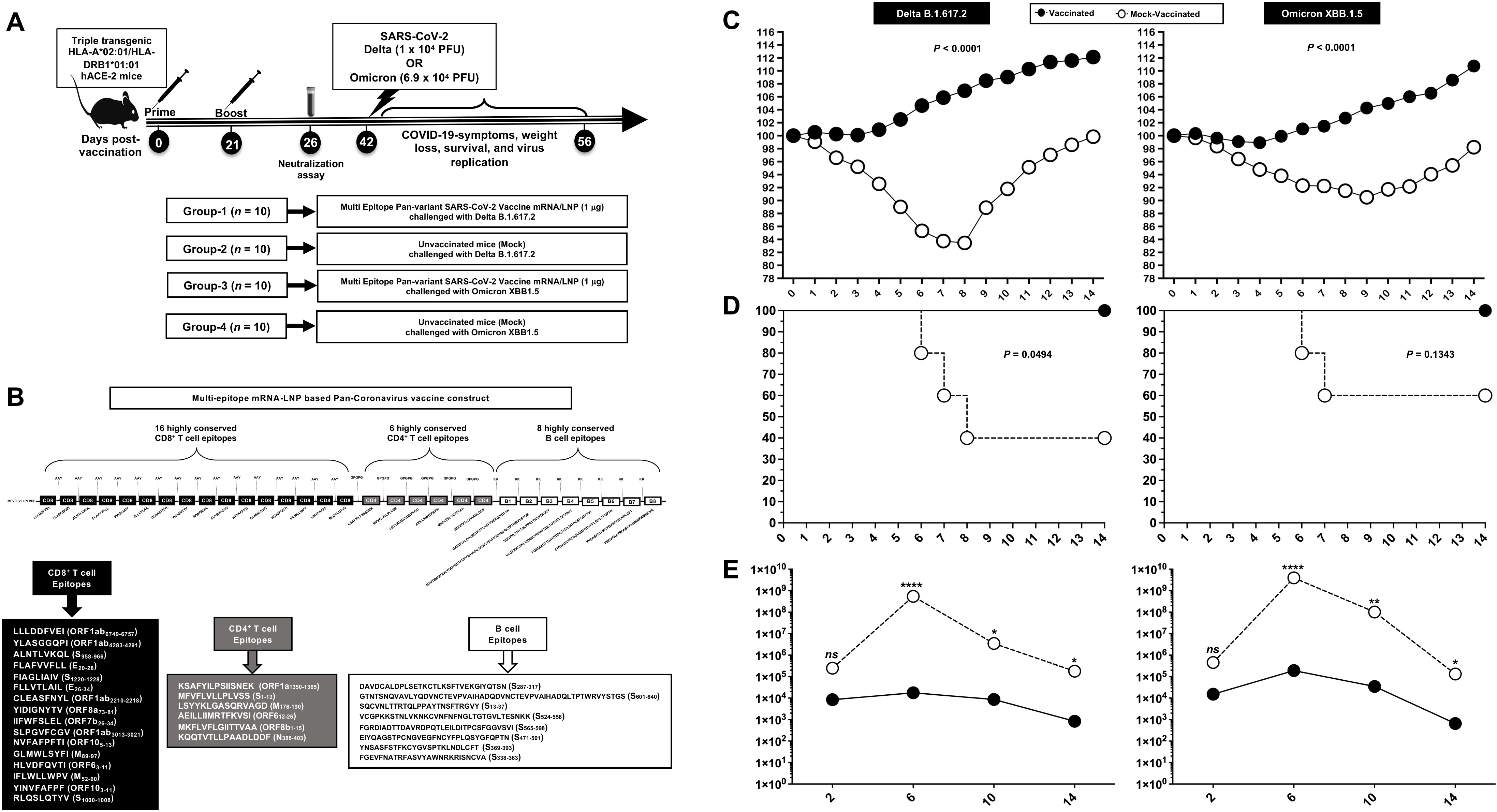
Protective efficacy of a mRNA-LNP based multi-epitope vaccine incorporating conserved human CD4^+^, CD8^+^ T, and B cell epitopes: (A) Experimental scheme of vaccination and challenge of triple transgenic HLA-A*02:01/HLA-DRB1*01:01-hACE-2 mice. Triple transgenic HLA-A*02:01/HLA-DRB1*01:01-hACE-2 mice (7-8-week-old, *n* = 20) were vaccinated intramuscularly (1 ug mRNA or mock) on Day 0 and Day 21 then challenged with Delta B.1.617.2 or Omicron XBB.1.5. (B) The B cell, CD4^+^ T cell, and CD8^+^ T cell epitope sequence details. (C) Percent weight change recorded daily for 14 days post-challenge in vaccinated and mock-vaccinated mice following the virus challenge. (D) Kaplan-Meir survival plots following virus challenge. (E) Virus replication following the virus challenge was detected in throat swabs on Days 2, 6, 10, and 14 post-challenge. The indicated *P* values are calculated using the unpaired *t*-test, comparing results obtained in vaccinated vs. mock-vaccinated mice.

### Vaccination and virus challenge

Animal studies were conducted under the University of California, Irvine IACUC Protocol number AUP-22-086 according to the NIH Guide for the Care and Use of Laboratory Animals. A sample size of 10 per group was determined to produce significant results with a power > 80%. Groups of 6- to 8-week-old female triple transgenic HLA-A*02:01/HLA-DRB1*01:01-hACE-2 mice were vaccinated intramuscularly at a 1 μg mRNA dose administered in 100 μl on days 0 and 21. Mock vaccinated controls received phosphate-buffered saline. On day 26, serum samples were collected for measurement of neutralizing antibodies. On day 42, mice were transferred to the ABSL-3 facility and intranasally challenged with the SARS-CoV-2 Delta variant (1 × 10^5^ pfu) or Omicron strain (2 × 10^5^ pfu).

### Virus titration

Oropharyngeal swabs were analyzed by qRT-PCR using ORF1ab-specific primers (For-ward-5′-CCCTGTGGGTTTTACACTTAA-3′ and Re-verse-5′-ACGATTGTGCATCAGCTGA-3′) and a probe (6FAM-CCGTCTGCGGTATGTGGAAAGGTTATGG-BHQ) to quantify SARS-CoV-2-specific RNA levels. Briefly, 100 ng of the total nucleic acid eluate was added to a 20 µL total-volume reaction mixture (1x TaqPath 1-Step RT-qPCR Master Mix (Thermo Fisher Scientific, Waltham, MA, USA), with 0.9 mM of each primer and 0.2 mM of each probe. qRT-PCR was carried out using the ABI StepOnePlus thermocycler (Life Technologies, Grand Island, NY, USA).

### Neutralizing antibodies

A serial dilution of 5% CO2 and 37°C was used to incubate SARS-CoV-2 variants with heat-inactivated plasma (1:3) for 30 minutes. Antibody-virus inoculums were neutralized by incubating at 34°C in 5% CO2, fixing in 10% neutral buffered formalin, incubating at −20°C for 10 minutes, then at room temperature for 20 minutes after neutralization. ELISpot reader was used to scan the plates developed with True Blue HRP substrate. IC50 was calculated based on normalized counted foci.

### Histology

To preserve mouse lungs, 10% neutral buffered formalin was used for 48 hours and then 70% ethanol was used for the next 24 hours. After embedding in paraffin blocks, tissue sections were sectioned at 8 mm thick. Routine immunopathology tests were performed after deparaffinization and rehydration of slides.

### Flow cytometry

Single-cell lung mononuclear cell suspensions were prepared by collagenase treatment (8mg/ml) for 1 hour and stimulated with peptide as described before ^10, 12, 14^. Cells were subsequently stimulated with 1µg/ml of each of the 16 CD8^+^ and 6 CD4^+^ individual T cell peptides and incubated in humidified 5% CO_2_ at 37°C. Subsequently, 1 x 10^6^ lung mononuclear cells (MNCs) were stained according to Dolton et al ^16^ for 45 minutes at room temperature with HLA-A*02*01 and/or HLA-DRB1*01:01 restricted tetramers (PE-labelled) specific toward Orf1ab_2210-2218_, Orf1ab_4283-4291_, S_1220-1228_, ORF10_3-11_ and toward the CD4^+^ T cell epitopes ORF1a_1350-1365_, S_1-13_, M_176-190_, ORF6_12-26_, respectively.

Surface staining was performed using anti-mouse CD4 (FITC, clone GK1.5 – BioLegend), CD8 (APC-Cy7, clone 53-6.7 – BioLegend), CD4^+^ Tetramer (APC), CD8^+^ Tetramer (PE), CD69 (BUV-496, clone H1.2F3 – BD), CXCR5 (BV786, clone L138D7 – BioLegend), and Ki-67 (BV-421, clone Ki-67 – BioLegend) in a total of 1 x 10^6^ cells in PBS containing 1% FBS and 0.1% Sodium azide for 45 minutes at 4°C. Cells were washed three times with FACS buffer and fixed in PBS containing 2% paraformaldehyde (Sigma-Aldrich, St. Louis, MO). A total of ∼100,000 lymphocyte-gated MNCs were acquired by Fortessa X20 (Becton Dickinson, Mountain View, CA) and analyzed using FlowJo software (Becton Dickinson, Ashland, OR). For the intracellular staining, after surface staining MNCs were treated with the Cytofix/Cytoperm (Becton Dickinson, USA), and were stained for 45 minutes at 4°C with Gzym-B (PerCP, clone QA16A02 – BioLegend). Cells were washed three times with FACS buffer and fixed in PBS containing 2% paraformaldehyde (Sigma-Aldrich, St. Louis, MO).

### Enzyme-linked immunosorbent assay (ELISA)

Assays were conducted on 96 well plates (Dynex Technologies, Chantilly, VA) coated with 100 ng spike protein or 0.5 grams of peptides overnight at 4°C, washed three times with PBS, and blocked with 3% BSA (in 0.1% PBST) for 2 hours at 37°C. A serial dilution of sera was added to the plates for 2 hours at 37°C after blocking. An HRP-conjugated goat anti-mouse IgG was used together with chromogenic substrate TMB (ThermoFisher, Waltham, MA) to detect bound serum antibodies.

## RESULTS

### A mRNA-LNP multi-epitope vaccine candidate, bearing conserved and asymptomatic B- and T-Cell epitopes protects against multiple SARS-CoV-2 variants of concern

We designed a multi-epitope mRNA-LNP vaccine encoding 16 conserved CD8^+^ T cell epitopes (ORF1ab_6749-6757_, ORF1ab_4283-4291_, S_958-966_, E_20-28_, S_1220-1228_, E_26-34_, ORF1ab_2210-2218_, ORF8a_73-81_, ORF7b_26-34_, ORF1ab_3013-3021_, ORF10_5-13_, M_89-97_, ORF6_3-11_, M_52-60_, ORF10_3-11_, and S_1000-1008_), 6 conserved CD4^+^ T cell epitopes (ORF1a_1350-1365_, S_1-13_, M_176-190_, ORF6_12-26_, ORF8b_1-15_, and N_388-403_), and 8 conserved B cell epitopes (S_1-13,_ S_287-317,_ S_13-17,_ S_524-558,_ S_471-501,_ S_369-393,_ S_338-363,_ and S_565-598_) (Figure 1B). These epitopes were selected from different structural and non-structural proteins of the SARS-CoV-2 genome ^12^ based on (1) their degree of conservation across diverse Coronavirus strains and (2) their selective and high recognition by B and T cells from unvaccinated asymptomatic COVID-19 patients, as reported previously. ^10^

Vaccine efficacy was evaluated in triple transgenic HLA-A*02:01/HLA-DRB1*01:01-hACE-2 mice to enable assessment of human T cell epitopes against challenge with SARS-CoV-2 Delta (B.617.2) and Omicron (XBB.1.5) variants, as measured by morbidity (weight loss), survival, virus replication, and lung pathology. The vaccine conferred complete protection from weight loss (Fig.1C) and death (Fig. 1D). In contrast, mock-vaccinated mice underwent substantial weight loss and death. Levels of virus replication, as measured in throat swabs, were significantly decreased (up to 5 logs) in vaccinated mice on days 6, 10, and 14 post-challenge for both Delta and Omicron variants (Fig. 1E). Lung pathology, as demonstrated by hematoxylin and eosin staining performed on lung sections at day 14 post-challenge, was markedly reduced in vaccinated versus mock vaccinated mice for both Delta (**Fig. 2A**) and Omicron (**Fig. 2B**) variants.

**Figure 2.**
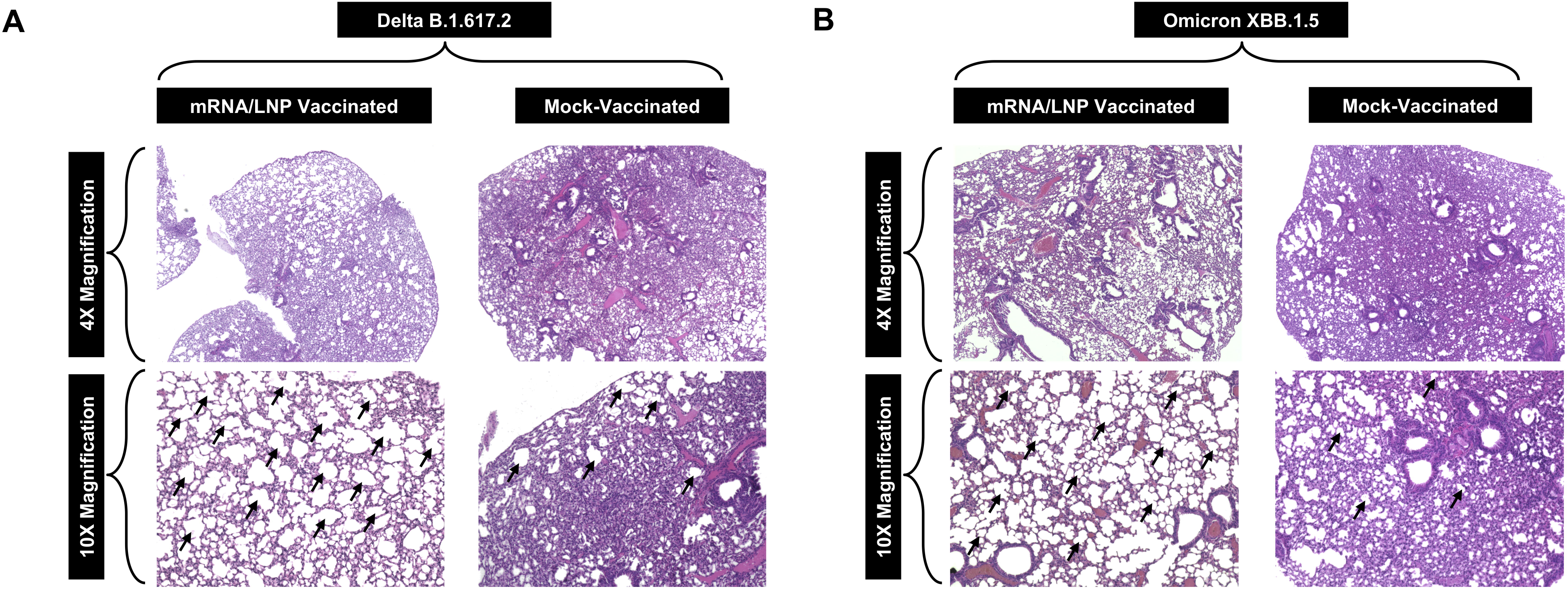
Lung histopathology in virus-challenged mice: (**A**) Representative images of hematoxylin and Eosin (H & E) staining of the lungs harvested on day 14 post-challenge from vaccinated (*left panels*) and mock-vaccinated (*right panels*) mice challenged with SARS-CoV-2 Delta (B.1.617.2) variant and (**B**) SARS-CoV-2 Omicron (XBB.1.5) variant. Black arrows point to the alveolar sacs. Images are shown at 4x, and 10x magnification.

Taken together, these results demonstrate that a multi-epitope mRNA-LNP vaccine based on carefully selected conserved and “asymptomatic” CD4^+^, CD8^+^ T, and B cell epitopes can confer substantial protection against infection and disease caused by SARS-CoV-2 variants.

### Potent neutralizing antibody responses are associated with protection induced by the mRNA-LNP multi-epitope vaccine candidate, bearing conserved and asymptomatic B- and T-Cell epitopes

The multi-epitope vaccine used herein contains eight conserved B cell epitopes selected from the SARS-CoV-2 Spike glycoprotein, three of which namely S_287-317_, S_524-558_, and S_565-598_ ^10, 12^ are completely conserved multiple variants (Supplementary Fig. 3). Neutralizing antibody responses were measured in sera collected on day 26 after first vaccination (i.e., 5 days after the second vaccination and before virus challenge) by a plaque reduction neutralization test. As shown in Fig. 3, the vaccine elicited antibodies that could neutralize both the Delta (B.1.617.2) and Omicron (XBB.1.5) viral variants.

### Frequent IFN-γ-producing CD8^+^ T cells in the lungs are induced by the mRNA-LNP multi-epitope vaccine candidate, bearing conserved and asymptomatic B- and T-Cell epitopes

The presence of IFN-γ-producing antigen-specific lung-resident CD8^+^ T cells was investigated in vaccinated and mock-vaccinated mice at 14 days post-challenge. Lung cell suspensions were stimulated with each of the 16 human CD8^+^ T cell epitopes described in *Materials & Methods* section and in Supplementary Fig. S1. ELISpot results showed a significant increase in the number of IFN-γ-producing CD8^+^ T cells in the lungs of vaccinated mice compared to mock-vaccinated mice irrespective of whether the epitopes were from structural (Spike, Envelope, Membrane) or non-structural (ORF1ab, ORF6, ORF7b, ORF10) SARS-CoV-2 protein antigens. This was true for mice challenged with either the Delta (Fig. 4A) or Omicron variant (Fig. 4C). Sequence homology analysis of the 16 human CD8^+^ T cell epitopes revealed complete conservation was maintained throughout the COVID-19 pandemic; as these 16 CD8^+^ T cell epitopes were 100% conserved against h-CoV-2/Wuhan, h-CoV-2/WA/USA2020, h-CoV-2/Delta (B.1.617.2), h-CoV-2/Omicron (BA.2), h-CoV-2/Omicron (XBB.1.5), h-CoV-2/Omicron (JN.1), h-CoV-2/Omicron (KP.2), and h-CoV-2/Omicron (KP.3) (Supplementary Fig. S1).

**Figure 3.**
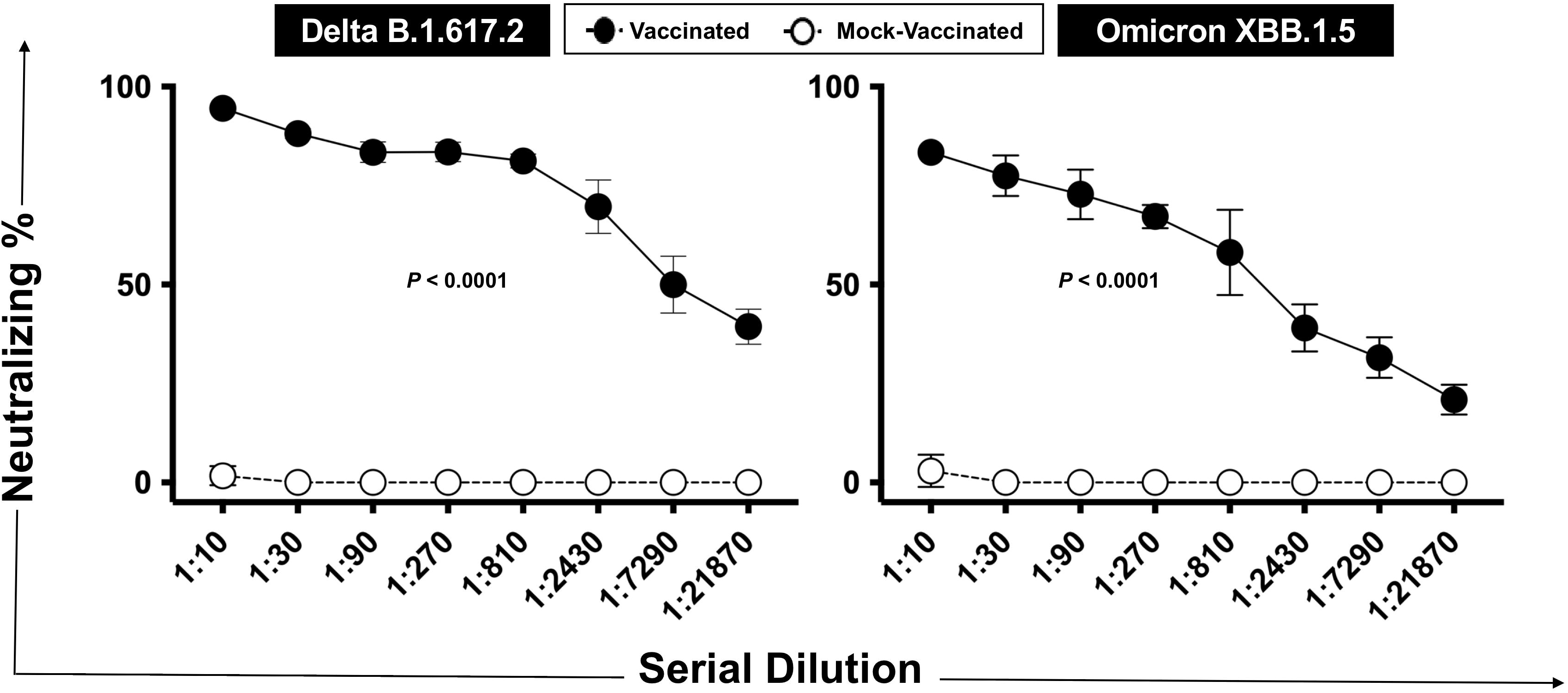
Neutralizing antibody responses: Serum samples were collected on Day 26 of the experiment (i.e., 5 days after the second immunization and before the challenge) from vaccinated and mock-vaccinated mice. Percent neutralization across 3-fold serum dilutions is shown for the two groups of mice destined to be challenged with the Delta and Omicron variants. Data were analyzed by student’s *t*-test and multiple t-tests. Results were considered statistically significant at *P* < 0.05. Statistical correction for multiple comparisons was applied using the Holm-Sidak method.

**Figure 4.**
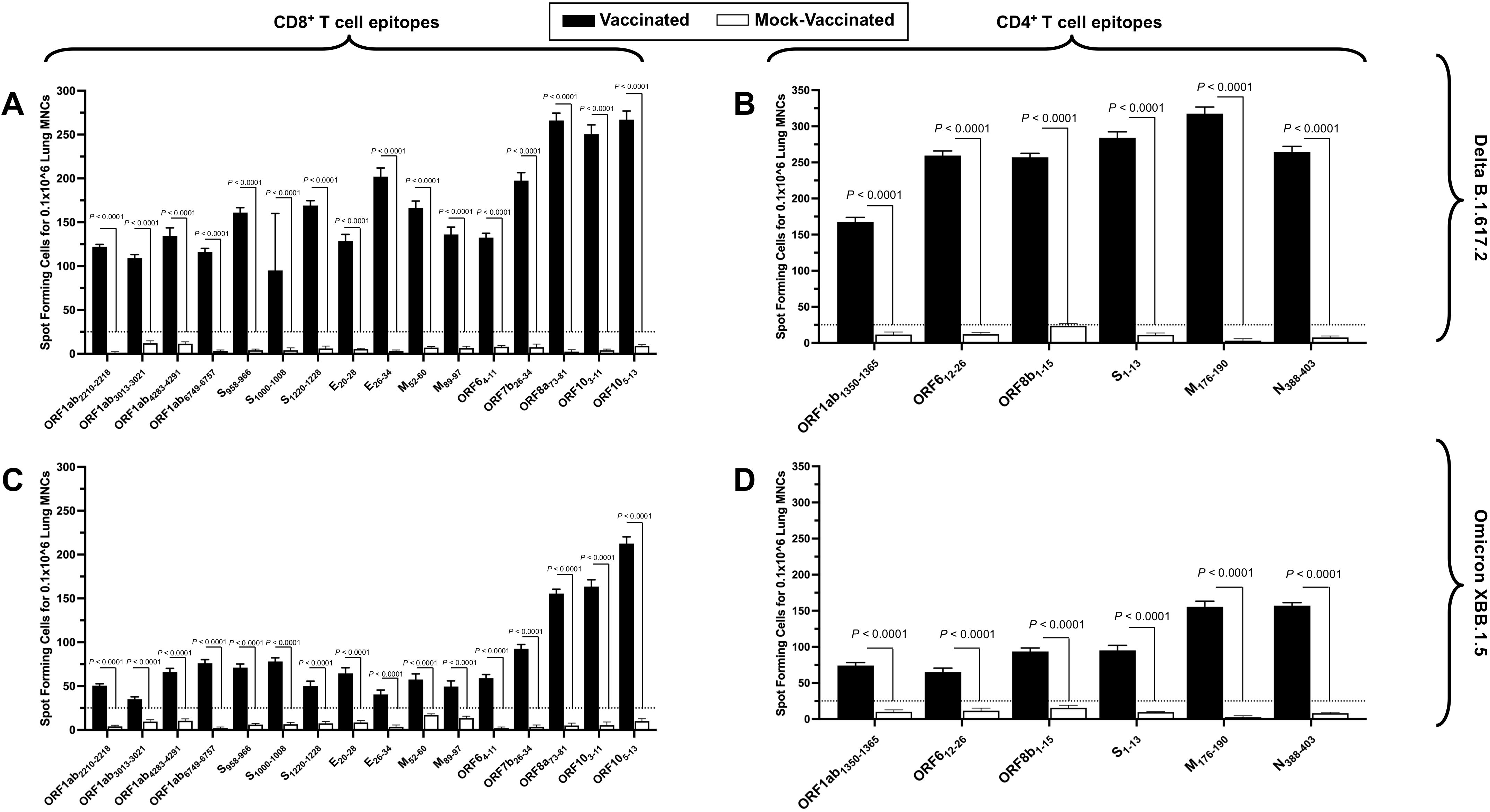
Induction of IFN-producing CD8^+^ T cell and CD4^+^ T cell responses: Shown are frequencies of IFN-producing cell ELISpots from lung mononuclear cells (1 x 10^6^ cells per well) of vaccinated and mock-vaccinated mice challenged with the Delta (B.1.617.2) (**A** & **B**) and Omicron (XBB.1.5) variants (**C** & **D**). CD8^+^ T cell responses are shown in panels A and C, while CD4^+^ T cells are shown in panels B and D. Dotted lines represent an arbitrary threshold set to evaluate the relative magnitude of the response. A strong response is defined for mean SFCs > 25 per 1 x 10^6^ stimulated MNCs. Results were considered statistically significant at *P* < 0.05.

### Frequent IFN-γ-producing CD4^+^ T cells in the lungs are induced by the mRNA-LNP multi-epitope vaccine candidate, bearing conserved and asymptomatic B- and T-Cell epitopes

The presence of of IFN-γ-producing antigen-specific lung-resident CD4^+^ T cells was investigated in vaccinated and mock-vaccinated mice at 14 days post challenge. Lung cell suspensions were stimulated with each of the 6 human CD4^+^ T cell epitopes described in *Materials & Methods* section and in Supplementary Fig. S2. ELISpot results showed significant increase in the number of IFN-γ-producing CD4^+^ T cells in the lungs of vaccinated mice compared to mock-vaccinated mice. Similar to the CD8+ T cell results, all comparisons showed statistically significant differences irrespective of whether the epitopes were from structural or non-structural protein antigens and whether the mice were challenged with the Delta (Fig. 4B) or Omicron variant (Fig. 4D). Sequence homology analysis of the 6 human CD4^+^ T cell epitopes demonstrated 100% conservation against all major variants of concern (Supplementary Fig. S2).

### Frequent functional CD8^+^ T cells in the lungs are induced by the mRNA-LNP multi-epitope vaccine candidate, bearing conserved and asymptomatic B- and T-Cell epitopes

To assess the function, activation and proliferation of antigen-specific lung-resident CD8^+^ T cells, the frequencies of lung-resident Gzym-B^+^ tetramer^+^ CD8^+^ T cells, CD69^+^ tetramer^+^ CD8^+^ T cells and Ki67^+^ tetramer^+^ CD8^+^ T against SARS CoV-2 Delta (B.1.617.2) and Omicron (XBB.1.5) variants were measured by flow cytometry. We observed significantly higher frequencies of tetramer^+^ CD8^+^ T cells for all 16 CD8^+^ T cell epitopes in the lungs of mice challenged with the Delta (Fig. 5A) and for all but 3 of the epitopes in the lungs of mice challenged with the Omicron variant (Fig. 6A). Gzym-B^+^ tetramer^+^ CD8^+^ T cells were found to be significant for 4 and 11 of the 16 CD8^+^ T cell epitopes in the lungs of mice challenged with the Delta (Fig. 5B) and Omicron (Fig. 6B) variants, respectively. With regard to the activation levels of epitope-specific lung-resident CD8^+^ T cells, significantly higher frequency of CD69^+^ tetramer^+^ CD8^+^ T cells were found for 10 and 4 of the 16 CD8^+^ T cell epitopes in the lungs of mice challenged with the Delta (Fig. 5C) and Omicron (Fig. 6C) variants, respectively. Finally, Ki67^+^ tetramer^+^ CD8^+^ T cells were used as a measure of proliferation status and we found higher frequencies for all 16 CD8^+^ T cell epitopes in the lungs of mice challenged with either the Delta (Fig. 5D) or Omicron (Fig. 6D) variants.

**Figure 5.**
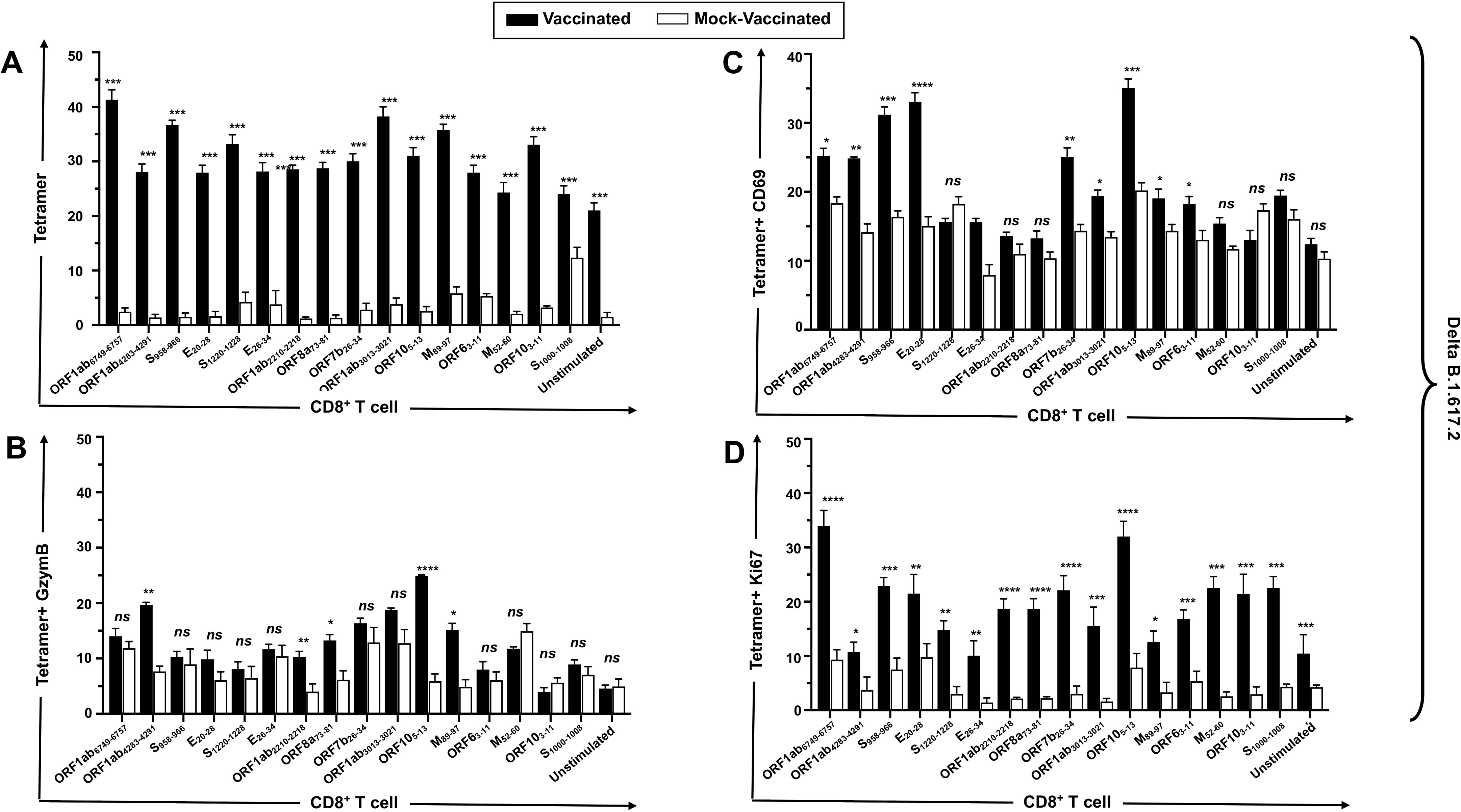
Induction of functional CD8^+^ T cells in mice challenged with the Delta variant: Frequencies of T cell responses after stimulation of lung cells with each 16 of the immunodominant CD8^+^ T cell peptides are shown for Tetramer^+^ CD8^+^ T cells (**A**), Tetramer^+^ IFN-γ^+^ CD8^+^ T cells (**B**), Tetramer^+^ CD69^+^ CD8^+^ T cells (**C**), and Tetramer^+^ Ki67^+^ CD8^+^ T cells (**D**). Bars represent means ± SEM. Data were analyzed by student’s *t*-test. Results were considered statistically significant at *P* < 0.05.

**Figure 6.**
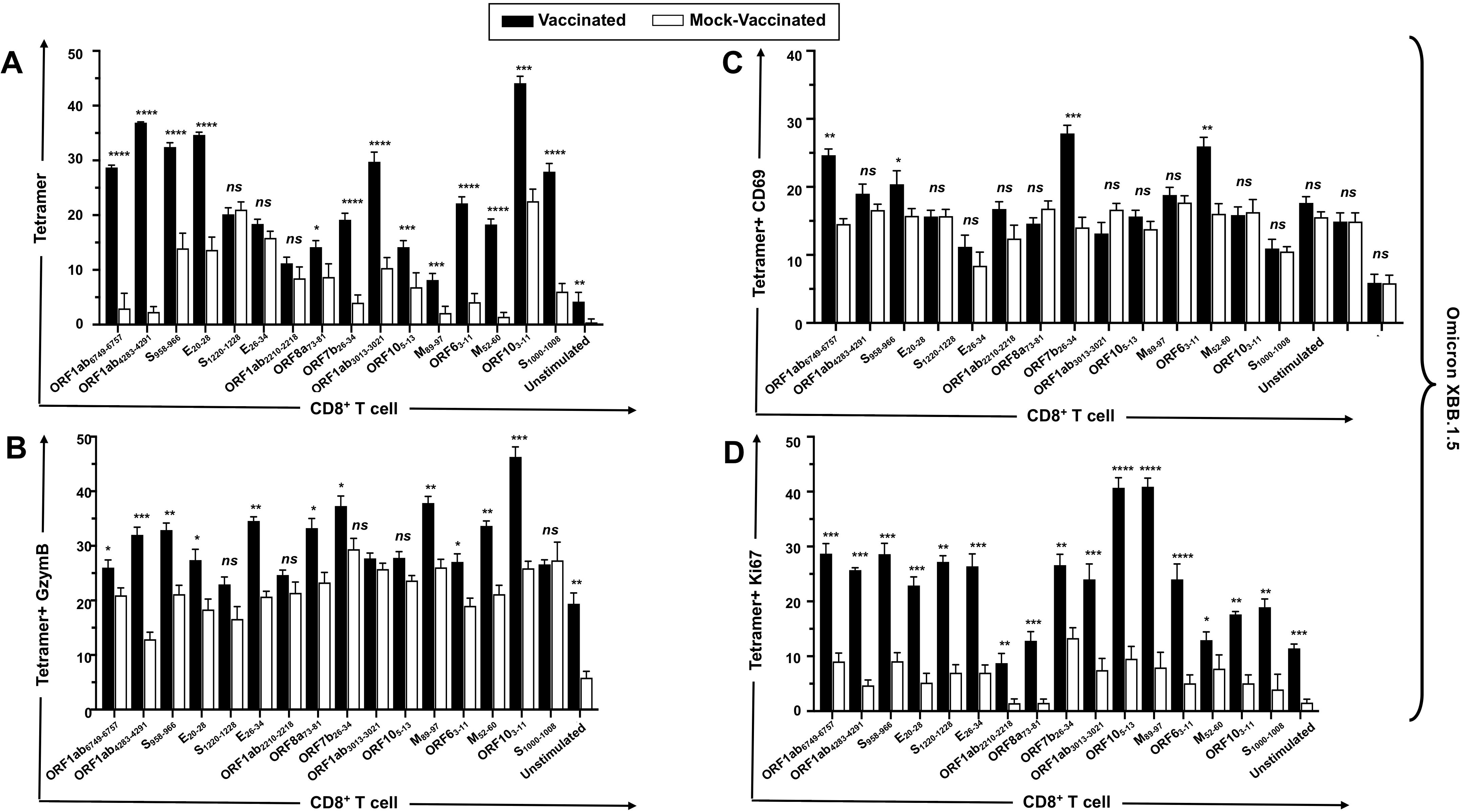
Induction of functional CD8^+^ T cells in mice challenged with the Omicron variant: Frequencies of T cell responses after stimulation of lung cells with each 16 of the immunodominant CD8^+^ T cell peptides are shown for Tetramer^+^ CD8^+^ T cells (**A**), Tetramer^+^ IFN-γ^+^ CD8^+^ T cells (**B**), Tetramer^+^ CD69^+^ CD8^+^ T cells (**C**), and Tetramer^+^ Ki67^+^ CD8^+^ T cells (**D**). Bars represent means ± SEM. Data were analyzed by student’s *t*-test. Results were considered statistically significant at *P* < 0.05.

### Frequent functional CD4^+^ T cells in the lungs are induced by the mRNA-LNP multi-epitope vaccine candidate, bearing conserved and asymptomatic B- and T-Cell epitopes

To determine whether the lung-resident tetramer-specific Follicular helper CD4^+^ T (T(FH)) cells were associated with protection in the vaccinated mice, lung cells were stimulated with each of the 6 human CD4^+^ T cell epitopes. Significantly higher frequencies of tetramer^+^ CD4^+^ T cells were observed for all 6 CD4^+^ T cell epitopes in the lungs of mice challenged with the Delta (Fig. 7A) and Omicron variants (Fig. 7E). Similarly, significantly higher frequencies of CXCR5^+^ tetramer^+^ CD4^+^ T cells were seen for 6 of 6 and 5 of 6 epitopes in the lungs of mice challenged with the Delta (Fig. 7B) and Omicron variants (Fig. 7F). With regard to the activation levels of lung-resident epitope-specific CD4^+^ T cells, significantly higher frequency of CD69^+^ tetramer^+^ CD8^+^ T cells were found for 6 and 5 of the 6 CD4^+^ T cell epitopes in the lungs of mice challenged with the Delta (Fig. 7C) and Omicron (Fig. 7G) variants, respectively. Finally, Ki67^+^ tetramer^+^ CD8^+^ T cells were used as a measure of proliferation status and we found higher frequencies for all 6 CD4^+^ T cell epitopes in the lungs of mice challenged with either the Delta (Fig. 7D) or Omicron (Fig. 7H) variants.

**Figure 7.**
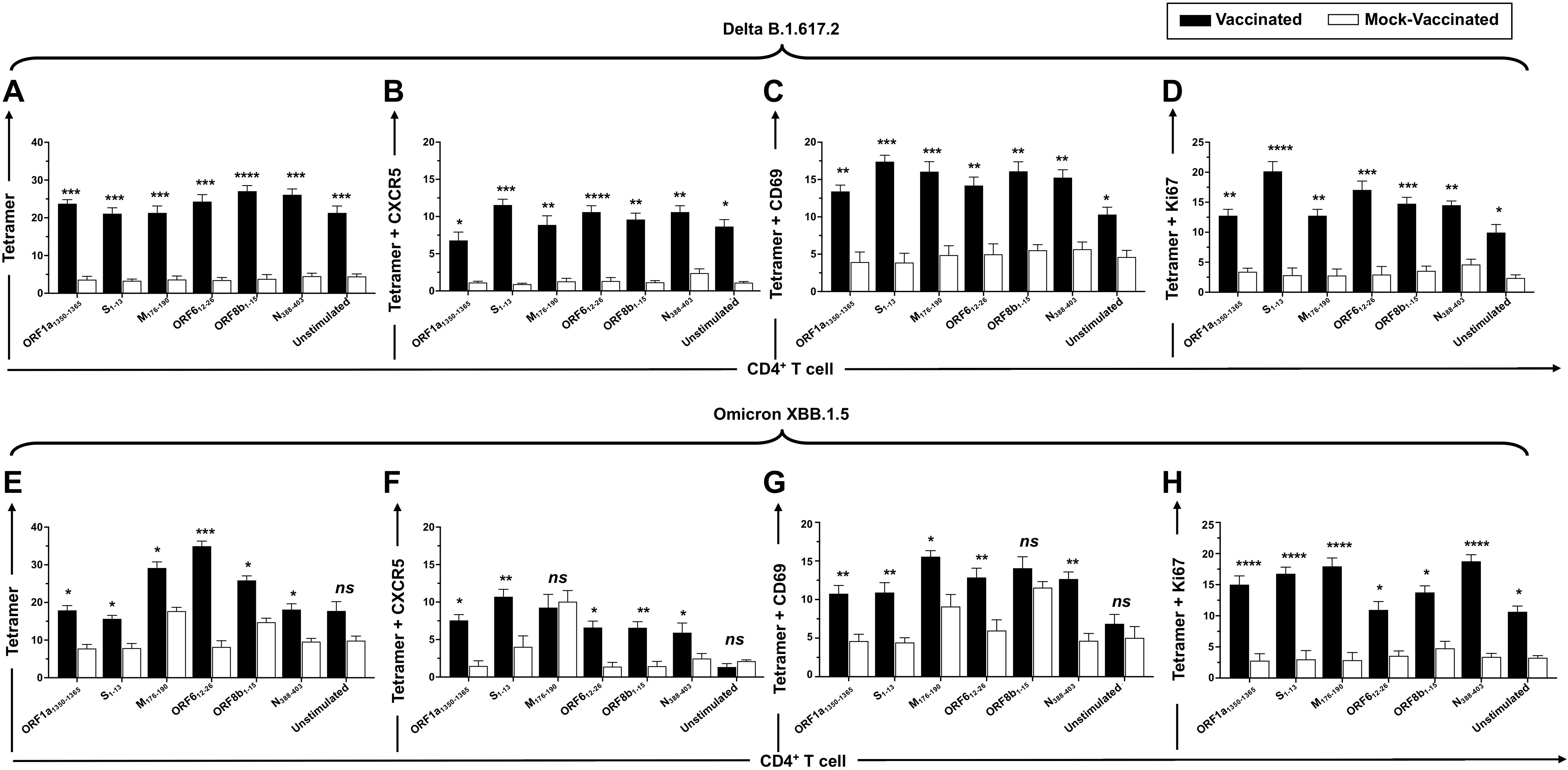
Induction of functional CD4^+^ T cells: Frequencies of T cell responses after stimulation of lung cells with each of the 6 immunodominant CD4^+^ T cell peptides are shown for Tetramer^+^ CD4^+^ T cells (A, E), Tetramer^+^ CXCR5^+^ CD4^+^ T cells (B, F), Tetramer^+^ CD69^+^ CD4^+^ T cells (C, G), and Tetramer^+^ Ki67^+^ CD4^+^ T cells (D, H) in mice challenged with the Delta (A-D) and Omicron variants (E-H). Numbers indicate frequencies of. Bars represent means ± SEM. Data were analyzed by student’s *t*-test. Results were considered statistically significant at *P* < 0.05.

In summary, these results demonstrate that a broad-spectrum mRNA-LNP multi-epitope vaccine, bearing conserved and “asymptomatic” B, CD4^+^, and CD8^+^ T cell epitope, elicited functional neutralizing antibodies and lung-resident CD4^+^ and CD8^+^ T cells that correlate with protection against challenge with the SARS-CoV-2 Delta and Omicron variants that are known to be highly pathogenic and transmissible.

## DISCUSSION

The COVID-19 pandemic, caused by SARS-CoV-2, has had a major global impact with deaths exceeding 7 million and economic damage in the trillions of dollars. Since 2020, there have been several distinct phases of the disease and humankind’s response to it. During the first year, intense efforts were marshaled worldwide with substantial investment to rapidly create effective vaccines. Remarkably, by the end of 2020, several new vaccines were introduced and rolled out, initially largely in the developed world. During the second year (2021), massive manufacturing scale-up and global distribution started to bring vaccines to most parts of the world. Yet, the disease was still killing up to ∼100,000 people per week. During the third year (2022), the trajectory of COVID-19 finally changed with mortality rates decreasing to ∼10,000 people per week, in part due to the widespread use of protective vaccines. However, while the vaccines reduced mortality and serious disease, they did not prevent infection and transmission of the virus from person to person. ^17^ Consequently, new virus variants appeared that were capable of evading vaccine-induced immunity resulting in the need for updated booster vaccines and a cyclical pattern of disease that has persisted ever since. In addition, the long-term consequences of COVID infection (i.e., Long COVID) can be devastating with lingering symptoms, including fatigue, breathing difficulties, cognitive issues (brain fog), muscle and joint pain, chest pain, and heart issues, observed in ∼10% of the global population with economic impacts reaching as high as $1 trillion annually. Fortunately, vaccination can reduce the chances of contracting Long COVID by ∼50%. While risk has declined over time due to vaccination and changes in the virus, unvaccinated populations remain disproportionally affected.

The current spike-based vaccines, while highly effective during the initial stages of the pandemic, have two key limitations due to the reliance on antibody responses directed toward a single viral antigen (i.e., spike). First, vaccine-induced antibody responses and immunity wane over time, potentially related to the paucity of long-lived antigen-specific plasma cells. ^18^ Second, SARS-CoV-2 evolution over time has produced a succession of variants containing a large number of mutations in the spike gene, thereby creating a mismatch between the vaccine and the circulating virus strain. As a consequence, vaccine-induced anti-spike antibodies are less effective at neutralizing these variants resulting in lower levels of protection. ^19^ Hence, improved vaccines providing broader and more durable immunity are needed.

Toward the goal of developing a broad-spectrum coronavirus vaccine, we have identified CD8^+^ and CD4^+^ T cell and B cell epitopes based on 1) conservation among all known SARS-CoV-2 variants, SARS-CoV-1 and MERS, and animal strains reported to be hosts (bat, civet cat, pangolin, camel), and 2) correlation of epitope-specific T cells responses with good disease outcomes in unvaccinated COVID-19 patients (). We designed and constructed a combination of these epitopes epitopes derived from both structural (Spike, Envelope, Membrane, Nucleocapsid) and non-structural (orf1ab, ORF6, ORF7, ORF8, and ORF10) proteins. To evaluate the protective efficacy of these human T cell epitopes, we developed a novel triple transgenic HLA-A*02:01/HLA-DRB1*01:01-hACE-2 mouse model and correlated these pre-clinical data with T cell reactivity in human COVID-19 subjects. The multi-epitope vaccine was demonstrated to provide robust protection against SARS-CoV-2 infection and COVID-19 disease caused by the Delta B.1.617.2 and Omicron XBB.1.5 variants, as measured by reduced morbidity (weight loss), higher survival rates, lower levels of virus replication, and improved lung pathology. Previously we observed that protection in the mouse model was associated with high frequencies of IFN-γ^+^ CD4^+^ T cells, CD69^+^ CD4^+^ T cells, IFN-γ^+^ CD8^+^ T cells, CD69^+^ CD8^+^ T cells, and TEM CD8^+^ (CD44^+^CD62L^-^) cells infiltrating the lungs ^12^ This was true irrespective of the viral variant used in the challenge. Similarly, in the present study, protection correlated with the presence of functional, antigen-specific T cells in the lungs. Specifically, higher levels of tetramer-specific GzymB^+^ CD8^+^ T cells, CD69^+^ CD8^+^ T cells, and Ki67^+^ CD8^+^ T cells were observed, relating to cytotoxicity, cellular activation, and proliferation, respectively. Higher levels of lung-resident tetramer-specific CXCR5^+^ CD4^+^ follicular helper T cells, CD69^+^ CD4^+^ T cells, and Ki67^+^ CD4^+^ T cells were also seen. The vaccine also elicited neutralizing antibody responses, likely due to the presence of B cell epitopes, which may have contributed to protection. Taken together, these data are supportive of targeting conserved B and T cell antigens to elicit broad-based immunity with the potential to confer cross-strain protection against a wide range of coronavirus variants.

The mRNA technology is well suited for the delivery of a multi-epitope vaccine targeting both humoral and cellular immunity. mRNA vaccines work by delivering a synthetic mRNA sequence encoding a viral antigen inside host cells, where the mRNA is translated and the ensuing antigen is presented to the immune system. This process mimics a virus infection and thereby triggers immune responses including the production of neutralizing antibodies and activation of T cells. ^20^ The mechanisms, efficacy, and immunogenicity of mRNA vaccines, including their role in combating SARS-CoV-2, are well described. ^4, 5^

Multi-epitope vaccines are designed to achieve broad-spectrum protection by targeting multiple epitopes targeting both B and T cell immunity. This is particularly significant for coronaviruses, which are prone to mutations in their spike protein. Hence, multi-epitope vaccines may reduce the risk of immune escape by targeting conserved regions across different viral strains. ^21^ Multiepitope vaccines can be designed to include (*i*) B cell epitopes to induce neutralizing antibodies, and (*ii*) T cell epitopes to activate cytotoxic T lymphocytes (CTLs) and follicular helper T cells, which are essential for clearing infected cells and providing long-term immune memory. ^21^ Comprehensive immune activation makes multi-epitope vaccines particularly attractive for use against complex pathogens such as coronaviruses as it has the potential to provide cross-protection against multiple coronavirus species and reduce the risk of future pandemics. Advances in bioinformatics and immune informatics have enabled the rapid identification of immunodominant and conserved epitopes. This allows for the quick design and development of multiepitope vaccines tailored to emerging pathogens. For example, computational tools can predict epitopes that are most likely to elicit strong immune responses, streamlining the vaccine development process. ^21^

Multi-epitope vaccines can be designed to exclude epitopes associated with adverse immune responses, such as those that trigger antibody-dependent enhancement (ADE) or excessive inflammation. This precision reduces the risk of vaccine-related side effects, making multi-epitope vaccines safer for widespread use. ^21^ Despite their promise, vaccines targeting discrete epitopes pose certain immunologic challenges, such as (*i*) identification of dominant human B and T cell epitopes, (ii) limitations in the number of epitopes that can be practically included may compromise coverage across the diversity of human leukocyte antigen types, and (iii) susceptibility to mutations in the pathogen that may reduce immune recognition of the vaccine epitope. However, ongoing research and technological advancements are expected to overcome these challenges, paving the way for the more widespread adoption of multi-epitope vaccines.

As the COVID-19 pandemic has entered its sixth year with no end in sight, vaccine research and development toward improved vaccines remains critical. Multi-epitope vaccines represent a significant advancement in vaccine technology, able to elicit targeted immunogenicity, confer broad-spectrum efficacy, and have the potential for long-term protection against rapidly evolving pathogens like coronaviruses. Their ability to target conserved and “asymptomatic” epitopes and elicit robust B- and T-cell responses makes them a promising tool in the fight against current and future pandemics.

## DISCLOSURE STATEMENT

LB has an equity interest in TechImmune, LLC., a company that may potentially benefit from the research results and serves on the company’s Scientific Advisory Board. LB’s relationship with TechImmune, LLC., has been reviewed and approved by the University of California, Irvine by its conflict-of-interest policies. Authors, HV, JU, and DG were employed by TechImmune, LLC. No potential conflict of interest was reported by the remaining authors.

## FUNDING

Studies of this report were supported by Public Health Service Research grants AI158060, AI150091, AI143348, AI147499, AI143326, AI138764, AI124911, and AI110902 from the National Institutes of Allergy and Infectious Diseases (NIAID) to LBM.

## DATA AVAILABILITY STATEMENT

The original contributions presented in the study are included in the article/supplementary material. Further inquiries can be directed to the corresponding author.

## ETHICAL APPROVAL AND CONSENT TO PARTICIPATE

The study is approved by UCI’s IBC approval number BUA-R112 and IACUC approval number AUP-22-086.

## INTRODUCTION OF THE CORRESPONDING AUTHOR

Dr.Lbachir BenMohamed is an immunologist and virologist who graduated from Pasteur Institute, in Paris, France, with a strong career focus on vaccine development for viruses. He has over 30 years of experience in viral infection/immunity and vaccine development, including a focus on coronavirus infection and immunity and on developing a pre-emptive pan-coronavirus vaccine, since early 2020. He has authored more than 120 peer-reviewed papers in immunology, virology, and vaccine development. Dr. BenMohamed’s team is recognized as a world leader in the fields of coronavirus and herpes T-cell immunity, memory T cells, and T-cell-based vaccines and immunotherapies. His group has pioneered “asymptomatic” viral CD4^+^ and CD8^+^ T cell epitope mappings for both herpes and coronaviruses and the identification of inflammasome pathways associated with inflammatory responses induced by virulent and non-virulent strains virus strains.

