## Supplemental Figure S1 for "A Pan-Beta-Coronavirus Vaccine Bearing Conserved and Asymptomatic B- and T-Cell Epitopes Protect Against Highly Pathogenic Delta and Highly Transmissible Omicron SARS-CoV-2 Variants of Concern"

Supplementary Fig. S1

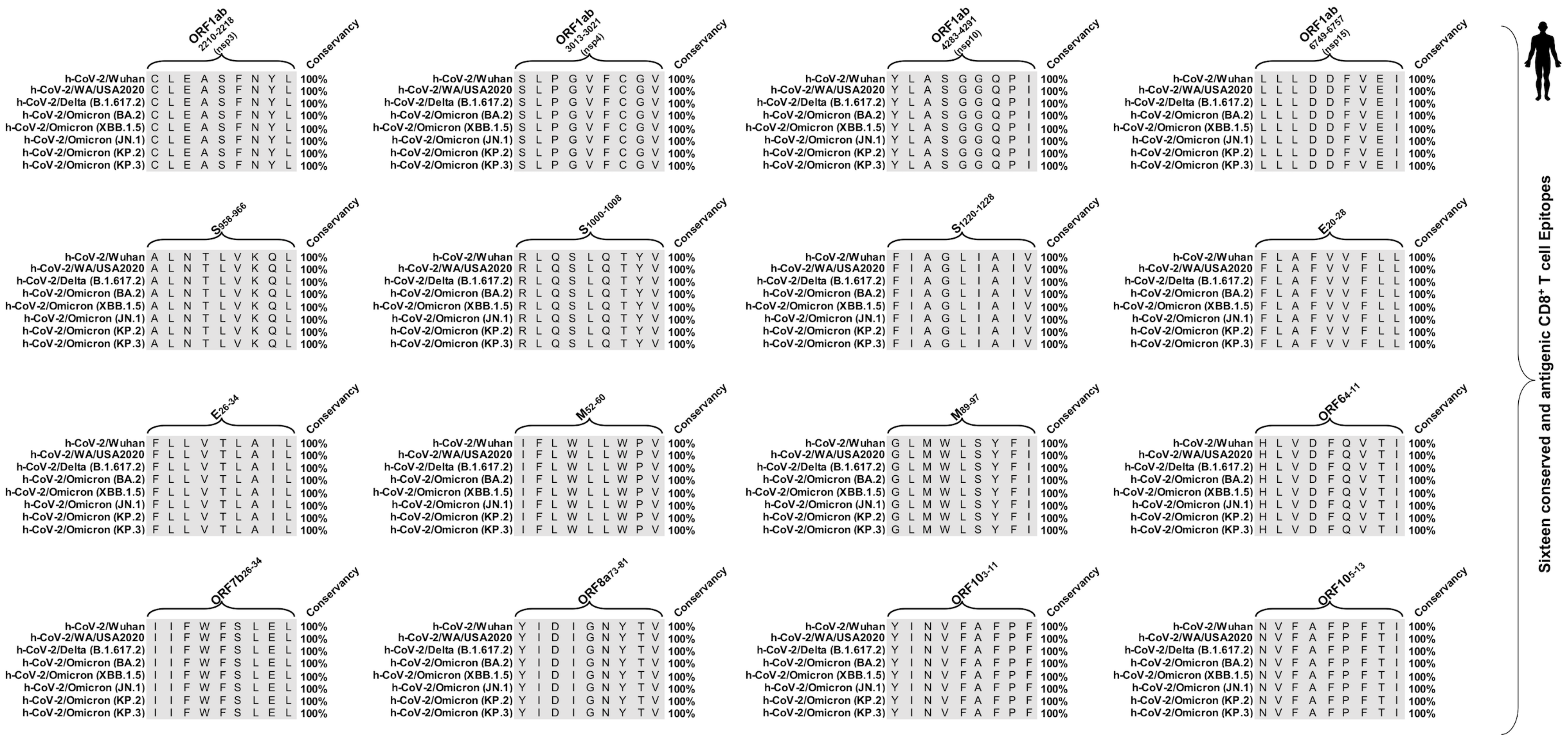


**Supplementary Figure S1. Sequence homology analysis to identify the degree of the conservancy of the immunodominant CD8$^{+}$ T cell epitopes among SARS-CoV-2 variants:** Sequence homology data for the CD8$^{+}$ T cell epitopes is shown. The 16 epitopes, found to be highly immunodominant against SARS-CoV-2 variants of concern WA/USA2020, Delta (B.1.617.2), Omicron (BA.2), Omicron (XBB.1.5), Omicron (JN.1), Omicron (KP.2), and Omicron (KP.3) were subjected to the sequence homology analysis.
