## Supplemental Figure S2 for "A Pan-Beta-Coronavirus Vaccine Bearing Conserved and Asymptomatic B- and T-Cell Epitopes Protect Against Highly Pathogenic Delta and Highly Transmissible Omicron SARS-CoV-2 Variants of Concern"

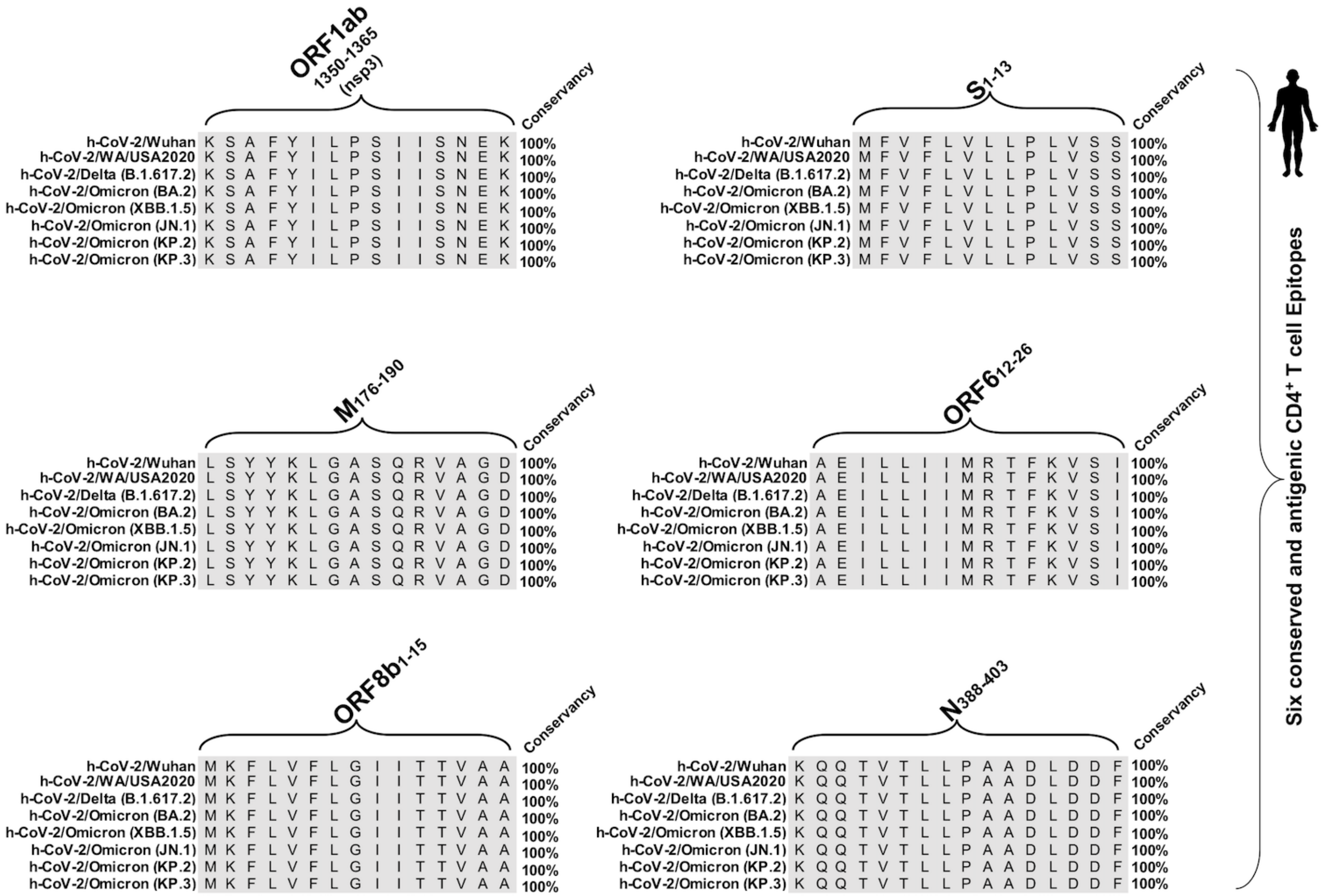


**Supplementary Figure S1. Sequence homology analysis to identify the degree of the conservancy of the immunodominant CD4+ T cell epitopes among SARS-CoV-2 variants:** Sequence homology data for the CD4+ T cell epitopes is shown. The 6 epitopes, found to be highly immunodominant against SARS-CoV-2 variants WA/USA2020, Delta (B.1.617.2), Omicron (BA.2), Omicron (XBB.1.5), Omicron (JN.1), Omicron (KP.2), and Omicron (KP.3) were subjected to the sequence homology analysis.
