## Supplemental Figure S2 for "A Pan-Beta-Coronavirus Vaccine Bearing Conserved and Asymptomatic B- and T-Cell Epitopes Protect Against Highly Pathogenic Delta and Highly Transmissible Omicron SARS-CoV-2 Variants of Concern"

*Supplementary Fig. S3*

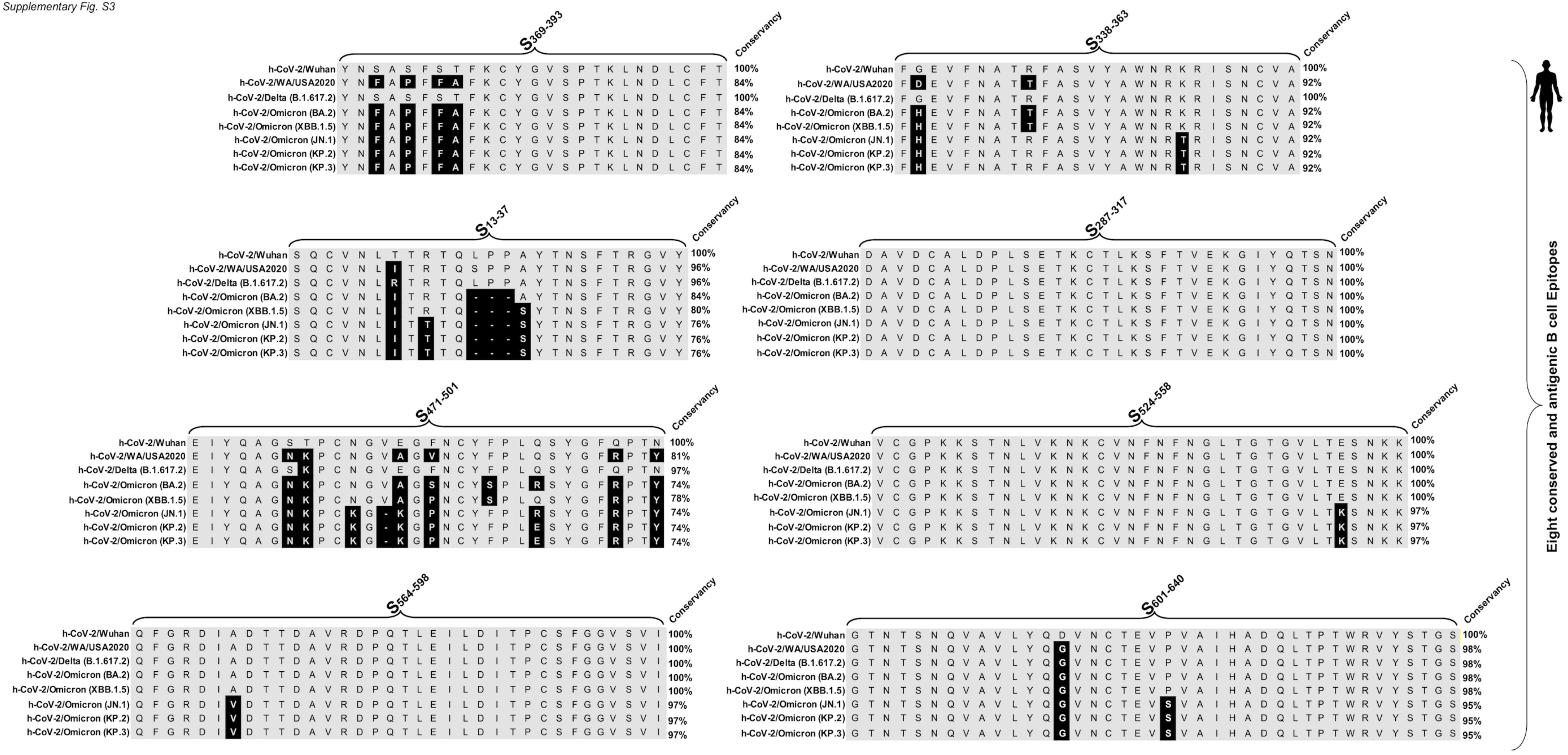


**Supplementary Figure S3. Sequence homology analysis to identify the degree of the conservancy of the immunodominant B cell epitopes among SARS-CoV-2 variants:** The sequence homology data for the B cell epitopes are shown. The 8 epitopes, found to be highly immunodominant against SARS-CoV-2 variants WA/USA2020, Delta (B.1.617.2), Omicron (BA.2), Omicron (XBB.1.5), Omicron (JN.1), Omicron (KP.2), and Omicron (KP.3) were subjected to the sequence homology analysis were subjected to the sequence homology analysis.
